# Is niche divergence more likely in parapatry? A test in *Sclerurus mexicanus* sensu lato (Aves: Furnariidae)

**DOI:** 10.1101/2022.12.06.519370

**Authors:** Jacob C. Cooper, Diego Barragán

**Author notes:** **Note**: This manuscript was written by the authors in 2015-2017, with this version approved for upload to Biorxiv in 2019, with minimal edits made in Dec 2022. This article was submitted and rejected pending revision; it is being posted here as a citable manuscript until the work can be revised appropriately and resubmitted. - JCC, 6 Dec 2022. Revision with proper GBIF DOI uploaded 12 Dec 2022.

## Abstract

**Aim:** While species’ niches are understood to diverge through time, the mode of these divergences is not fully understood. The null hypothesis of niche divergence in the present is that of conservatism; that species are likely to occur in the same environment as their ancestors and, therefore, as allopatric sister species. Alternatively, species are expected to diverge when selective pressure, such as parapatry with a congener, occurs. Here, we analyse niche divergence in a mosaic of allo- and parapatric *Sclerurus mexicanus* sensu lato populations to determine if niche divergence is more likely in parapatry.

**Location:** Neotropics.

**Methods:** We created a dataset of 1,100 vetted localities for *S. mexicanus* sensu lato, and assigned each point to each of the seven described populations. We created individual dispersal areas for each species in Maxent following: 1) a broad approach in which training areas where allowed to overlap with parapatric congeners; and 2) a narrow approach in which species dispersal areas adhered to strict parapatry and there was no overlap in training areas. We complemented these ENMs with ‘random’ models drawn from points within each training area, thus creating ‘null’ ENMs against which we could test niche divergence using the metric Schoener’s D.

**Results:** There was no significant difference in the performance of broad and narrow training areas. Few significant divergences were found, and all of those that were found consisted of allopatric populations. Partial divergences were frequently recovered when comparing parapatric taxa.

**Main conclusions:** In contrast to our expectation, we found no significant evidence for increased niche divergence in parapatric versus allopatric species. Possible explanations for these findings include true conservatism even among parapatric lineages (*e.g*., additional environmental or biological factors may delimit distributions) or artefactual errors inherent to model creation. We discuss the implications of these findings, and discuss ways to improve upon tests of niche divergence in the future, especially when dealing with cases of parapatry.

## INTRODUCTION

Since the conception of ecological niche theory, attempts have been made to quantify the differences between the occupied niche space of different taxa (Grinnell, 1917; Elton, 1927; Hutchinson, 1957; Colwell & Futuyma, 1971; Carnes & Slade, 1982; Austin et al., 1990; Glor & Warren, 2011). These differences have been hypothesized to be key or causal to species’ diversification (Orr & Smith, 1998), especially in cases of related species’ groups that inhabit spatially parapatric niches. In recent years, new methods have been put forth to test how species’ niches differ in ecological space, all utilizing different approaches to Ecological Niche Modeling (ENM), and all with the overarching goal of testing whether or not species truly inhabit different multivariate environmental (*i.e*., Hutchinsonian) niches (Hutchinson, 1957; Warren et al., 2008; Blonder et al., 2014; Nunes & Pearson, 2016).

As our knowledge of species’ diversification has increased, it has become apparent that most species diverge in allopatry by means of genetic drift, differential selection, and a lack of genetic mixing (Price, 2008). Likewise, our expectation of niche divergence has approached the null; that is, when species become allopatric, we hypothesize that they are more likely retain their ancestral niche and therefore appear ‘conserved’ via ENM tests (Peterson et al., 1999; McCormack et al., 2009; Peterson, 2011; Ryan Shipley et al., 2013; Khaliq et al., 2015; Skeels & Cardillo, 2017). Furthermore, research has shown that competition and the re-establishment of contact can drive niche (and character) divergence, and thereby result in the para- and sympatric shifts we observe in the present (Jang & Gerhardt, 2006; Kirschel et al., 2009; Wooten & Gibbs, 2012; Freeman, 2015; Miller et al., 2016).

While ecological niche divergence has been examined in allopatric complexes (e.g., McCormack et al., 2009), it has been less studied in parapatric species complexes. Given the expectation in allopatric speciation that niches remain conserved until secondary contact with a competitor occurs, it is therefore logical to hypothesize that the magnitude of niche divergence will be greater when species have a contact zone during their speciation cycle (Nunes & Pearson, 2016). This hypothesis is reinforced by our understanding of niche shifts that accompany speciation events involving species moving into parapatric habitats or geographic realms (Benkman et al., 2009; Fjeldså et al., 2011; Knowles & Massatto, 2016), and has provided the basis of a novel methodology for testing ecological niche divergence (Nunes & Pearson, 2016). There are few tests of the assumption of niche divergence within monophyletic parapatric groups (Reino et al., 2016), and it is difficult to fully assess the other factors that may be influencing species’ realized niches in ENMs.

Soberón & Peterson (2005)elucidate ways to address additional factors in their **BAM** framework, highlighting three factors that are responsible for limiting species’ distributions: (**B**) biotic interactions; (**A**) abiotic environmental conditions (*i.e*., scenopoetic variables *sensu* Soberón, 2007); and (**M**) motility restrictions, such as geographic barriers (*i.e*., an invasible area). Biotic interactions, collectively known as a species’ Eltonian niche (Soberón, 2007), are often assumed to be implicitly accounted for in ENMs by a study species’ absence from areas with a biotic amensal or without a biotic commensal. Linking these factors directly into an ENM is often done by restricting a species’ explorable area in the initial steps of ENM creation, thereby removing regions of potential sympatry from model training (Saupe et al., 2012; de Araújo et al., 2014; Cooper & Soberón, 2018). ENMs can also opt to ignore these biotic factors, as many of the localized competitors and commensals a species encounters may not be as important as abiotic factors in large-scale spatial analyses (*i.e*., the Eltonian Noise Hypothesis; Soberón, 2007; Soberón & Nakamura, 2009).

In order to better understand niche divergence in parapatry, we performed a multidirectional niche-difference test of a widespread species group, *Sclerurus mexicanus* sensu lato (Cooper & Barragán, 2016). Now known to comprise of at least five major groups of taxa (d’Horta et al., 2013; Cooper & Cuervo, 2017), populations of *S. mexicanus* occur in allo- and parapatric populations from central Mexico to southern Brazil, and exhibit remarkable levels of ecomorphic and phenotypic conservatism, with field reports of species’ responding to unrelated, allopatric population’s songs (Remsen, 2003; Cooper & Barragán, 2016). Given their lack of known sympatry, it is likely that parapatric populations are competitors with other members of the super species, and therefore provide an excellent mosaic of parapatric and allopatric populations in which to test the magnitude of niche divergence in the presence and absence of amensals.

Here, we perform two sets of niche divergence analyses on this species complex under two alternative hypotheses: 1) that species can freely disperse into regions occupied by parapatric congeners, and 2) that species are prohibited from dispersal into regions where congeners are known occur. While both of these methods will model a species’ absence in areas with direct competitors, their mechanisms in reaching these conclusions differ. The first hypothesis excludes environments that are more optimal for competitive congeners by accounting for these as “absent” environments for the species of interest, while the second hypothesis manually thresholds environments and excludes these areas from analyses altogether. The first hypothesis also assumes that species have differential competitive abilities in different parts of their shared niche space or differ to at least some extent in their fundamental niches, while the second hypothesis assumes that competitive exclusion is masking the limits of a species’ ecological niche. While there is no way to get around these errors for attempting to assess a species’ complete fundamental niche, it is still possible to analyze a species’ realized niche and determine whether or not species are significantly differing in their occupied ecological space. Furthermore, a test of both methods will help reveal whether biotic exclusion is significantly affecting our interpretations of ecological niche divergence.

## METHODS

We downloaded occurrence data for *Sclerurus mexicanus* sensu lato from the Global Biodiversity Informatics Facility (GBIF.org, 2015) and from eBird (EBD_relMay-2015; Sullivan et al., 2009; eBird, 2012). These data were combined with a set of georeferenced recordings from Cooper & Cuervo (2017), and were subsequently separated into species groups corroborated by the recordings. We recognized five species within the complex sensu Cooper & Cuervo (2017), but separated subspecies of *S. macconnelli* for niche analyses in light of the genetic monophyly of *peruvianus* with respect to *macconnelli* and the uncertain relationships of *bahiae* (d’Horta et al., 2013). We thus modeled seven recognized populations from *S. mexicanus* sensu lato, ordered in this manuscript from north to south and listed here with the number of vetted occurrences used for each: *Sclerurus mexicanus* (*n* = 532); *S. pullus* (*n* = 358); *S. obscurior* (*n* = 19); *S. andinus* (*n* = 24); *S. macconnelli peruvianus* (*n* = 77); *S. macconnelli macconnelli* (*n* = 41); and *S. macconnelli bahiae* (*n =* 49; Fig. 1). Localities were crossreferenced with the literature, and we omitted records that were unidentifiable by range or lacked physical documentation, most notably: intermediate elevations (*c*. 1200-1800 m) in the western Andes of Colombia and Ecuador where *S. obscurior* and *S. andinus* potentially overlap; an unidentified population from intermediate elevations from southern Ecuador; and an unidentified population from Ceará, Brazil (Cooper & Barragán, 2016).

**Figure 1.** The estimated geographic distribution of different *Sclerurus mexicanus* sensu lato populations from and basemap from Cooper & Cuervo (2017) with an overlay of training points from this study. Question marks indicate areas in which the population’s identity has not been corroborated with vocal data or genetic data.

Based on these occurrence data, we created accessible areas in which to train the models (hereafter, **M**s, sensu Soberón & Peterson, 2005 and Cooper & Soberón, 2018) using QGIS 2.8.2 (QGIS Development Team, 2016). **M**s were bounded by major biogeographic barriers (*e.g*., mountain ridgelines, major rivers, ecoregion boundaries [World Wildlife Fund, 2010], etc.) and incorporated what is hypothesized to be the entire accessible area for the species (Soberón & Peterson, 2005; Guisan et al., 2006). We created two types of **M**: broad **M**s that encompassed the geographic distribution of a species and included regions of potential dispersal, including areas in which a parapatric species occurs (*i.e*., hypothesis 1); and narrow **M**s that excluded the entirety of the distribution of a parapatric species from the training area (and are thus a subset of the broad **M**s), with divisions approximated halfway between known populations (*i.e*., hypothesis 2; Fig. 2).

**Figure 2.**
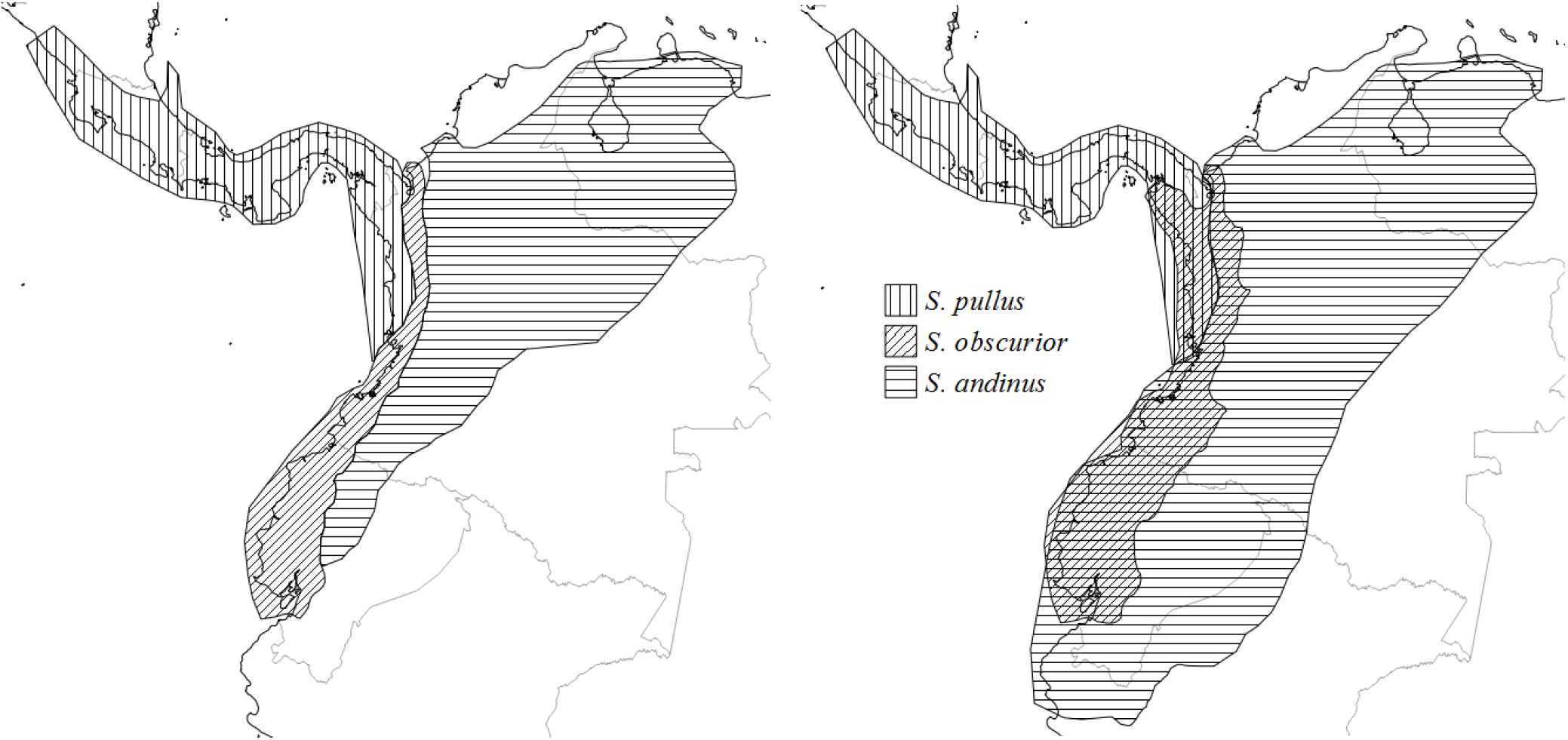
Narrow (left) and broad (right) training areas for three study taxa, listed from north to south: *S. pullus, S. obscurior*, and *S. andinus*. In the narrow training areas, strict parapatry (*i.e*., no sympatry) is enforced.

Analyses were performed in R 3.2.5 (R Core Team, 2016), with our study following the methodology of Warren et al. (2008). Bioclimatic data were downloaded at a resolution of 2.5 arcminutes (*c*. 5 km at the equator) for four bioclimatic variables: mean temperature of the warmest quarter, mean temperature of the coldest quarter, mean precipitation of the wettest quarter, and mean precipitation of the driest quarter (Hijmans et al., 2005). Suitability models were created using the average of five raw, unclamped outputs of MaxEnt 3.3.3k (Phillips et al., 2004), using the packages ‘dismo’ (Hijmans et al., 2015), ‘raster’ (Hijmans, 2015), and ‘rJava’ (Urbanek, 2013). Suitability models were projected into a larger ‘clade’ area (an **M** encompassing the entire super species) so that all populations could be modeled within each others’ distributions and suitability comparisons could be made in identical spatial and environmental extents.

Sets of random points were drawn from each population’s appropriate **M** to create environmentally ‘null’ suitability predictions in MaxEnt for contrast against actual population comparisons using the R package ‘maptools’ (Bivand & Lewin-Koh, 2015). These random point sets are equal in number to the known occurrences and exclude exact coordinates of known occurrences. We used these points to create null ENMs within the **M** of each species for each hypothesis using the same methodology outlined for individual species’ models. Following Warren et al. (2008), we used the metric of Schoener’s D to assess niche similarity, where the metric *D* is an indicator of the similarity between MaxEnt outputs, with total dissimilarity being 0 and identical outputs being 1. We replicated these analyses 100 times bi-directionally for each of the 21 pairwise comparisons for each hypothesis to create two independent random distributions of Schoener’s D using the R package ‘dismo’ (Warren et al., 2008; Glor & Warren, 2011; Peterson et al., 2011; Hijmans et al., 2015). If the Schoener’s D value of the direct comparison was less than the critical value (i.e. within the lower 5%) of both pseudoreplicate distributions, the comparison was scored as a “2” for being significantly divergent and the null hypothesis of niche conservatism was rejected; if the value was significant with regard to only one distribution, the comparison was ranked as “1” for being potentially divergent, but the results were not considered a rejection of overall niche conservatism; no differences between niches was ranked as “0”, and indicated a failure to reject the null of niche conservatism. Plots for all comparisons were visually inspected using the R package ‘ggplot2’ (Wickham, 2009). We did not assess extreme conservatism, as this is indistinguishable from the null hypothesis for ecological niches between allopatric populations. Differences in performance between the two methods was assessed via a Fisher’s Exact Test on the number of successes in allopatry and parapatry for broad and narrow **M**s and on the sum of the net amount of divergence for these categories.

## RESULTS

A total of 21 pair-wise comparisons of *Sclerurus mexicanus* sensu lato populations for both the broad and the narrow **M**s were completed (Table 1). Using broad training areas, three of these comparisons showed significant divergence (i.e. a “2”) among the populations compared, with all three of these involving comparisons with *S. macconnelli bahiae*. Twelve comparisons did not reject the null hypothesis of niche conservatism, but indicated potential unidirectional (slight) divergence (i.e. a “1”) while the remaining six comparisons showed no evidence for ecological niche divergence of any kind. In almost every case of potential divergence, the critical value fell within the range of the values derived from a montane population’s model compared to pseudoreplicates of its lowland congener, and not vice versa (Fig 3).

**Figure 3.**
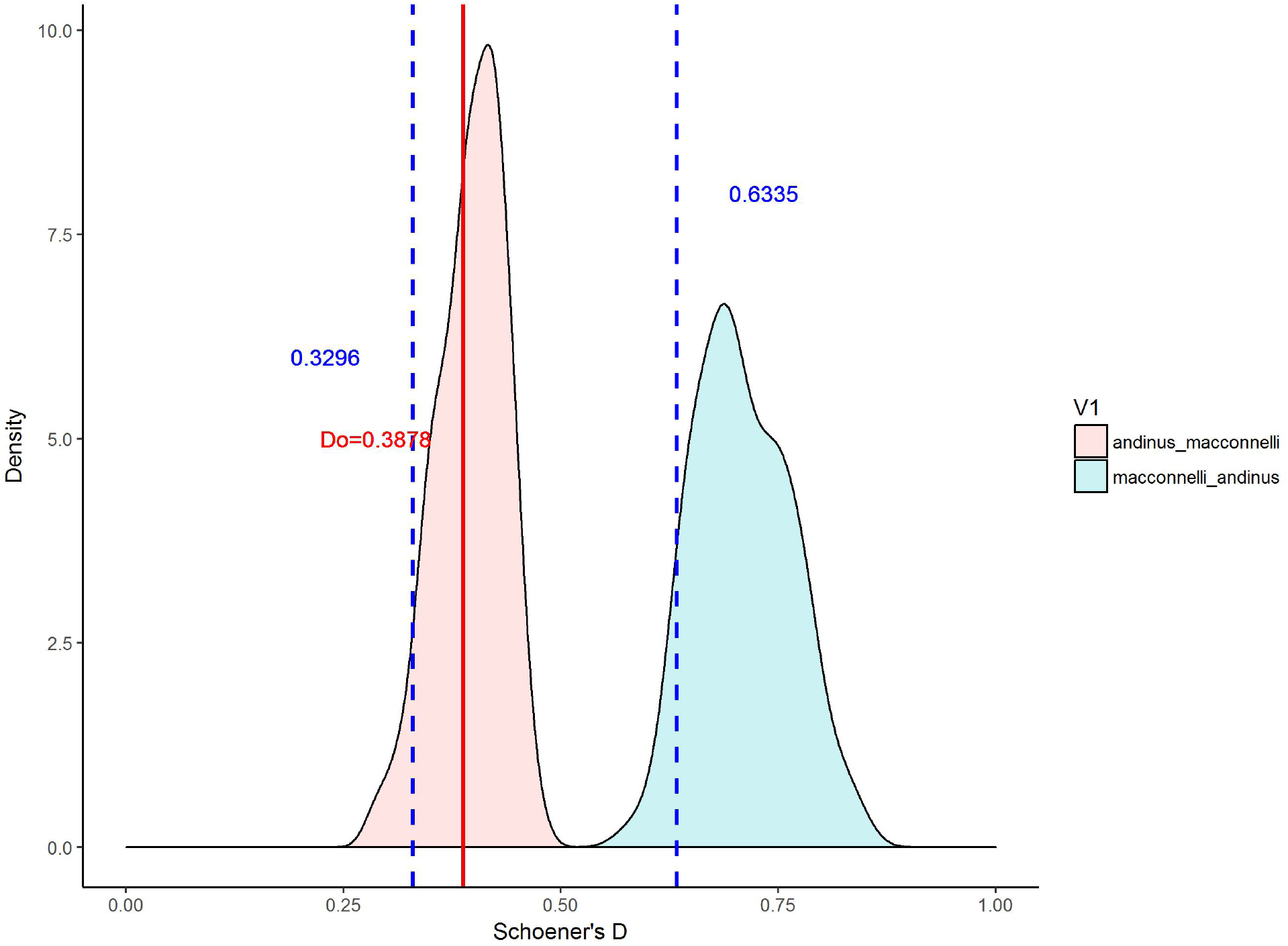
A comparison of broad models for *Sclerurus andinus* and *Sclerurus m. macconnelli*. The red (left) distribution is the random iterations of *S. andinus* compared to the true *S. m. macconnelli*, while the blue-green (right) distribution is the random iterations of *S. m. macconnelli* compared to *S. andinus*. Vertical dashed lines represent the critical values for this test (i.e. the point at which any test to the left has a *P* value of 0.05 or lower), while the solid vertical is the test statistic of the true ENM of *S. andinus* compared to the true ENM of *S. m. macconnelli*. As with many tests of montane vs. lowland taxa, the test was significant for the distribution of random lowland derived models, but not for random montane models.

**Table 1.** Pairwise comparisons of Schoener’s D for all combinations of *S. mexicanus* populations. Significance is ranked thusly: 2 = comparison was significant for the tails of both bidirectional comparisons; 1 = test was significant for one of the bidirectional comparisons; 0 = test was not significant.

| Populations | Broad *M* Results | Narrow *M* Results | Allopatric<br>? | Parapatric? |
| --- | --- | --- | --- | --- |
| *andinus-bahiae* | 2 | 1 | Yes |  |
| *andinus-macconnelli* | 1 | 1 | Yes |  |
| *andinus-mexicanus* | 0 | 0 | Yes |  |
| *andinus-obscurior* | 1 | 1 |  | Yes |
| *andinus-peruvianus* | 0 | 0 |  | Yes |
| *andinus-pullus* | 1 | 0 | Yes |  |
| *bahiae-macconnelli* | 0 | 1 | Yes |  |
| *bahiae-mexicanus* | 2 | 2 | Yes |  |
| *bahiae-obscurior* | 1 | 1 | Yes |  |
| *bahiae-peruvianus* | 1 | 0 | Yes |  |
| *bahiae-pullus* | 2 | 0 | Yes |  |
| *macconnelli-mexicanus* | 1 | 1 | Yes |  |
| *macconnelli-obscurior* | 1 | 0 | Yes |  |
| *macconnelli-peruvianus* | 1 | 1 |  | Yes |
| *macconnelli-pullus* | 1 | 1 | Yes |  |
| *mexicanus-obscurior* | 1 | 1 | Yes |  |
| *mexicanus-peruvianus* | 0 | 0 | Yes |  |
| *mexicanus-pullus* | 0 | 1 | Yes |  |
| *obscurior-peruvianus* | 1 | 1 | Yes |  |
| *obscurior-pullus* | 1 | 1 |  | Yes |
| *peruvianus-pullus* | 0 | 1 | Yes |  |

When species were modeled in narrow regions, the number of highly significant tests dropped, with only one “2” result obtained (*S. mexicanus* vs. *S. macconnelli bahiae*). In total, five comparisons lost value (e.g., decreased from 1 to 0), eleven remained the same, and three gained value (always from 0 to 1), with no significant difference in **M** performance. There was no observable trend as to whether allo- or parapatry increased the likelihood of obtaining a significant result with regards to niche divergence, nor was there any evidence that particular combination of habitat increased the probability of significance changing.

## DISCUSSION

Evidence for niche divergence was strongest (i.e., a 2) only in comparisons of the Atlantic forest *S. macconnelli bahiae* to the montane and semi-montane *S. mexicanus, S. pullus*, and *S. andinus*. Both **M** scenarios appeared to perform equally well, with the greatest divergences restricted to broad models between aforementioned allopatric populations. These results confirm the null hypothesis that niches will remain largely conserved through time, and that niche divergence in allopatric populations only occurs when coupled with differential selective pressures. In our tests, we failed to find strong support for increased niche divergence in parapatric versus allopatric taxa, despite the expectation that secondary contact drives species into different environments.

Despite a lack of evidence of increased divergence in parapatric taxa, it is worth noting that almost every instance of potentially diverging niches was the result of a direct comparison falling within the Schoener’s D distribution derived from comparing the montane population to random points from a lowland population. Given that *Sclerurus* as a whole is predominately a lowland genus, these results may indicate competitive exclusion of highland taxa from lowland areas. Alternatively, these results may indicate elevational stacking of species upon secondary contact where species segregate based on the elevations at which they outcompete one another. Such a scenario would result in different realized niches, but we would be unable to recover significant differences via ENM tests as the environmental niches would be abutting and potentially overlapping. Field tests of populations’ physiological tolerances and habitat preferences will be needed to determine whether there are uni- or bidirectional amensal relationships dictating elevational turnover within *Sclerurus mexicanus* sensu lato.

Conversely, these skewed divergences may be an artefact of model design, wherein montane habitats are usually proportionately smaller than adjacent lowland habitats and the probability of selecting montane points for a null model is thus lower. Thus, lowland taxa whose **M**s abut mountain ranges are more likely to have random points drawn from lowlands, whereas montane taxa are also more likely to have lower elevational limits drawn in a random process. Narrow models, while limiting the amount of habitat at different elevations that can be drawn, fail to account for potential biotic interactions between related taxa and assume *a priori* that taxa that are not explicitly sympatric are amensal. Further studies into the best ways to mitigate inherently unequal areas of habitat between species with different elevational ranges are needed to understand the best way to compensate for these inequalities, and the usage of presence-only techniques may be warranted (*e.g*., minimum volume ellipsoids; Van Aelst & Rousseeuw, 2009).

We are aware of one other study that assessed multiple related taxa in a mosaic of allo- and parapatric distributions: McCormack et al. (2009). They likewise failed to find strong support for niche divergence in allopatric populations, and noted the apparent conservatism of niches within their study genus, *Aphelocoma* (Aves: Corvidae). McCormack et al. (2009) used defined distributions to determine areas for drawing random points and filtering occurrence data, but apparently did not use **M**s to constrain taxa-specific ENMs. Similar to this study, many of the taxa in question are restricted to montane habitats and thus have a predilection for random points being drawn from lowland habitats, especially when dispersal limitations are not enforced for individual ENMs. Coincidentally, McCormack et al.’s (2009) sympatric species pair possessed Schoener’s D distributions closely resembling our parapatric comparisons, wherein the test indicates unidirectional niche divergence.

### Implications

Given the results of this and other studies using Warren et al.’s (2008) methodology for determining niche divergence, it is possible that parapatric species may not be recoverable as possessing different ecological niches using this framework. We know from specimen and vocal data that members of *Sclerurus mexicanus* sensu lato occupy elevational gradients in the Andes, and thus infer that these species must possess some differences in their ecological niches. However, the topography of these regions and the nature of the modeling programs used may make quantifying full niche divergence impossible. Furthermore, the potential biotic element of these scenarios – amensal relationships with congeners – implies that species’ distributions may be extremely proximate in nature and thus difficult to fully distinguish using broad-scale ENM procedures. The absence of a competitor in part of a species’ distribution may also influence the recovered environmental niche breadth; *i.e*., species may overlap environmentally with competitors in models due to records from areas where competitors are not present and thus not limiting a species’ niche or distribution (Connell, 1961).

Investigations into presence-only methods for determining niche divergence are needed, as well as continued explorations of lesser-used variables such as elevation and tree cover (Cord et al., 2014; Title & Bemmels, 2017). While differences in fundamental niche will necessitate field research into species’ physiological tolerances, presence-only methods will serve to circumvent topographical and areal biases that are introduced from pseudoabsence data. Further examinations of niche difference tests are also needed to determine if potential unidirectional divergences are typical for proximate species that differ in their realized ecological niches. While the results we present are indicative of widespread niche conservatism, it is entirely plausible that these taxa still exhibit some realized niche divergence with respect to some other ecological factor, and further investigation is warranted.

## ACKNOWLEDGMENTS

We are grateful to the following individuals and groups for their input on this project: John M. Bates, The Field Museum; Andrés Cuervo, Universidad de Los Andes; Chris Hensz, A. Townsend Peterson, & Jorge Soberón, University of Kansas Biodiversity Institute; the ENM group of the University of Kansas; and the ornithology group of The Field Museum. This work was part of JCC’s Masters research funded by NSF 1208472 and by a University of Kansas summer fellowship, and was part of DB’s undergraduate research at the same institution.

## LITERATURE CITED

Van Aelst S. & Rousseeuw P. (2009) Minimum volume ellipsoid. Wiley Interdisciplinary Reviews: Computational Statistics, 1, 71–82.

de Araújo C.B., Marcondes-Machado L.O., & Costa G.C. (2014) The importance of biotic interactions in species distribution models: a test of the Eltonian noise hypothesis using parrots. Journal of Biogeography, 41, 513–523.

Austin M.P., Nicholls A.O., & Margules C.R. (1990) Measurement of the Realized Qualitative Niche: Environmental Niches of Five Eucalyptus Species. Ecological Monographs, 60, 161–177.

Benkman C.W., Smith J.W., Keenan P.C., Parchman T.L., & Santisteban L. (2009) A New Species of the Red Crossbill (Fringillidae: *Loxia*) from Idaho. The Condor, 111, 169–176.

Bivand R. & Lewin-Koh N. (2015) maptools: Tools for Reading and Handling Spatial Objects. R package version 0.8-36.

Blonder B., Lamanna C., Violle C., & Enquist B.J. (2014) The n-dimensional hypervolume. Global Ecology and Biogeography, 23, 595–609.

Carnes B.A. & Slade N.A. (1982) Some Comments on Niche Analysis in Canonical Space. Ecology, 63, 888–893.

Colwell R.K. & Futuyma D.J. (1971) On the Measurement of Niche Breadth and Overlap. Ecology, 52, 567–576.

Connell J.H. (1961) The influence of interspecific competition and other factors on the distribution of the barnacle *Chthamalus stellatus*. Ecology, 42, 710–723.

Cooper J.C. & Barragán D. (2016) Tawny-throated Leaftosser (*Sclerurus mexicanus*). Neotropical Birds Online (T. S. Schulenberg, Editor).

Cooper J.C. & Cuervo A.M. (2017) Vocal variation and species limits in the *Sclerurus mexicanus* complex. The Wilson Journal of Ornithology, 129, 13–24.

Cooper J.C. & Soberón J. (2018) Creating individual accessible area hypotheses improves stacked species distribution model performance. Global Ecology and Biogeography, 27, 156–165.

Cord A.F., Klein D., Gernandt D.S., Pérez de la Rosa J.A., & Dech S. (2014) Remote sensing data can improve predictions of species richness by stacked species distribution models: a case study for Mexican pines. Journal of Biogeography, 41, 736–748.

d’Horta F.M., Cuervo A.M., Ribas C.C., Brumfield R.T., & Miyaki C.Y. (2013) Phylogeny and comparative phylogeography of *Sclerurus* (Aves: Furnariidae) reveal constant and cryptic diversification in an old radiation of rain forest understorey specialists. Journal of Biogeography, 40, 37–49.

eBird (2012) eBird: An online database of bird distribution and abundance [web application].

Elton C.S. (1927) Animal Ecology. Sidgwick and Jackson, London, United Kingdom,

Fjeldså J., Bowie R.C.K., & Rahbek C. (2011) The Role of Mountain Ranges in the Diversification of Birds. Annual Review of Ecology, Evolution, and Systematics, 43, 249–265.

Freeman B.G. (2015) Competitive interactions upon secondary contact drive elevational divergence in tropical birds. The American Naturalist, 186, 470–479.

GBIF.org. (2015) GBIF Occurrence Download https://doi.org/10.15468/dl.39f5dl

Glor R.E. & Warren D. (2011) Testing ecological explanations for biogeographical boundaries. Evolution, 65, 673–683.

Grinnell J. (1917) The niche-relationships of the California Thrasher. The Auk, 34, 427–433.

Guisan A., Lehmann A., Ferrier S., Austin M., Overton J.M., Aspinall R., & Hastie T. (2006) Making better biogeographical predictions of species’ distributions. Journal of Applied Ecology, 43, 386–392.

Hijmans R.J. (2015) raster: Geographic Data Analysis and Modeling. R package version 2.3-40.

Hijmans R.J., Cameron S.E., Parra J.L., Jones P.G., & Jarvis A. (2005) Very high resolution interpolated climate surfaces for global land areas. International Journal of Climatology, 25, 1965–1978.

Hijmans R.J., Phillips S.J., Leathwick J., & Elith J. (2015) dismo: Species Distribution Modeling.

Hutchinson G.E. (1957) Concluding Remarks. Cold Springs Harbor Symposia on Quantitative Biology, 22, 415–427.

Jang Y. & Gerhardt H.C. (2006) Divergence in the calling song between sympatric and allopatric populations of the southern wood cricket *Gryllus fultoni* (Orthoptera: Gryllidae). Journal of Evolutionary Biology, 19, 459–472.

Khaliq I., Fritz S.A., Prinzinger R., Pfenninger M., Böhning-Gaese K., & Hof C. (2015) Global variation in thermal physiology of birds and mammals: evidence for phylogenetic niche conservatism only in the tropics. Journal of Biogeography, 42, 2187–2196.

Kirschel A.N.G., Blumstein D.T., & Smith T.B. (2009) Character displacement of song and morphology in African tinkerbirds. Proceedings of the National Academy of Sciences, 106, 8256–8261.

Knowles L.L. & Massatto R. (2017) Distributional shifts - not geographic isolation - as a probable driver of montane species divergence. Ecography, 40, 1475–1485.

McCormack J.E., Zellmer A.J., & Knowles L.L. (2009) Does Niche Divergence Accompany Allopatric Divergence in Aphelocoma Jays As Predicted Under Ecological Speciation?: Insights From Tests With Niche Models. Evolution, 65, 1–14.

Miller E.T., Wagner S.K., Harmon L.J., & Ricklefs R.E. (2016) Radiating despite a Lack of Character: Ecological Divergence among Closely Related, Morphologically Similar Honeyeaters (Aves: Meliphagidae) Co-occurring in Arid Australian Environments. The American Naturalist, 189, E000–E000.

Nunes L.A. & Pearson R.G. (2016) A null biogeographic test for assessing ecological niche evolution. Journal of Biogeography, 1–7.

Orr M.R. & Smith T.B. (1998) Ecology and speciation. Trends in Ecology & Evolution, 13, 502–6.

Peterson A.T. (2011) Ecological niche conservatism: a time-structured review of evidence. Journal of Biogeography, 38, 817–827.

Peterson A.T., Soberón J., Pearson R.G., Anderson R.P., Martínez-Meyer E., Nakamura M., & Araújo M.B. (2011) Ecological niches and geographic distributions. Princeton University Press, Princeton, New Jersey.

Peterson A.T., Soberón J., & Sanchez-Cordero V. (1999) Conservatism of ecological niches in evolutionary time. Science, 285, 1265–1267.

Phillips S.J., Dudík M., & Schapire R.E. (2004) A maximum entropy approach to species distribution modeling. Proceedings of the Twenty-First International Conference on Machine Learning, 655–662.

Price T.D. (2008) Speciation in Birds. Roberts & Company Publishers, Greenwood Village, Colorado.

QGIS Development Team (2016) QGIS Geographic Information System. Open Source Geospatial Foundation Project,.

R Core Team (2016) R: A Language and Environment for Statistical Computing. R Foundation for Statistical Computing, Vienna, Au.

Reino L., Ferreira M., Martínez-Solano Í., Seguardo P., Xu C., & Barbosa A.M. (2017) Favourable areas for co-occurrence of parapatric species: niche conservatism and niche divergence in Iberian tree frogs and midwife toads. Journal of Biogeography, 44, 88–98.

Remsen J.V.J. (2003) Family Furnariidae (ovenbirds). Handbook of the Birds of the World: Vol. 8. Broadbills to Tapaculos (ed. by J. del Hoyo, A. Elliott, and D.A. Christie), pp. 162–357. Lynx Ediciones, Barcelona.

Ryan Shipley J., Contina A., Batbayar N., Bridge E.S., Peterson A.T., & Kelly J.F. (2013) Niche conservatism and disjunct populations. The Auk, 130, 476–486.

Saupe E.E., Barve V., Myers C.E., Soberón J., Barve N., Hensz C.M., Peterson A.T., Owens H.L., & Lira-Noriega A. (2012) Variation in niche and distribution model performance: The need for a priori assessment of key causal factors. Ecological Modelling, 237–238, 11–22.

Skeels A. & Cardillo M. (2017) Environmental niche conservatism explains the accumulation of species richness in Mediterranean-hotspot plant genera. Evolution, 71, 582–594.

Soberón J. (2007) Grinnellian and Eltonian niches and geographic distributions of species. Ecology Letters, 10, 1115–1123.

Soberón J. & Nakamura M. (2009) Niches and distributional areas: Concepts, methods, and assumptions. Proceedings of the National Academy of Sciences, 106, 19644–19650.

Soberón J. & Peterson A.T. (2005) Interpretation of Models of Fundamental Ecological Niches and Species’ Distributional Areas. Biodiversity Informatics, 2, 1–10.

Sullivan B.L., Wood C.L., Iliff M.J., Bonney R.E., Fink D., & Kelling S. (2009) eBird: An online database of bird distribution and abundance [web application]. Biological Conservation, 142, 2282–2292.

Title P.O. & Bemmels J.B. (2018) ENVIREM: An expanded set of bioclimatic and topographic variables increases flexibility and improves performance of ecological niche modeling. Ecography, 41, 291–307.

Urbanek S. (2013) rJava: Low-level R to Java interface. R package version 0.9-6.

Warren D.L., Glor R.E., & Turelli M. (2008) Environmental niche equivalency versus conservatism: Quantitative approaches to niche evolution. Evolution, 62, 2868–2883.

Wickham H. (2009) ggplot2: Elegant Graphics for Data Analysis. Springer-Verlag, New York.

Wooten J.A. & Gibbs H.L. (2012) Niche divergence and lineage diversification among closely related Sistrurus rattlesnakes. Journal of Evolutionary Biology, 25, 317–328.

World Wildlife Fund (2010) Terrestrial ecoregions. List of ecoregions.

